# Computer vision and deep learning automates nocturnal rainforest ant tracking to provide insight into behavior and disease risk

**DOI:** 10.1101/454207

**Authors:** Natalie Imirzian, Yizhe Zhang, Christoph Kurze, Raquel G. Loreto, Danny Z. Chen, David P. Hughes

## Abstract

Determining how ant colonies optimize foraging while mitigating disease risk provides insight into how the ants have achieved ecological success. Fungal infected cadavers surround the main foraging trails of the carpenter ant *Camponotus rufipes*, offering a system to study how foragers behave given the persistent occurrence of disease threats. Studies on social insect foraging behavior typically require many hours of human labor due to the high density of individuals. To overcome this, we developed deep learning based computer vision algorithms to track foraging ants, frame-by-frame, from video footage. We found foragers can be divided into behavioral categories based on how straight they walk across the trail. Eighty percent of ants walk directly across the trail, while 20% wander or circle when crossing the trail. Departure from the main trail encourages exploration of new areas and could enhance discovery of new food resources. Conversely, results from our agent-based model simulations suggest deviation from a straight path exposes foragers to more infectious fungal spores. Consistency in walking behavior may protect most ants from infection, while the foragers with increased exposure due to their mode of walking could be a sufficient number of new hosts to sustain disease in this environment.

## Introduction

Resource acquisition drives animals into new territories, while threat avoidance limits where animals move. A consistent threat is the presence of infectious propagules of parasites and these are hypothesized to be major determinants of the distribution of animals in the wild^1^. Examples of animals avoiding pathogen contaminated areas span diverse taxa, from mammals to insects, implying anti-parasite behavior is widespread^1–5^. Central place foragers are interesting in the context of parasite avoidance as they must obtain food while avoiding threats with the additional constraint of returning to a defined location after each trip. For volant central place foragers, like wasps, bees, bats and birds, much of the trip is through the air likely reducing contact with infectious material. However, for taxa which walk on the ground (e.g. ants), encounters with parasite propagules are presumably higher^6^. To effectively study such pressure, it is crucial to use systems where we can study foragers in nature, surrounded by their naturally occurring pathogens.

The foraging strategies of ants range from workers searching and retrieving food entirely independently to obligately in a group^7^. Chemical trails commonly facilitate group foraging, and in some cases, these chemical trails develop into semi-permanent trails known as ‘trunk trails’^8^. Trunk trails stimulate research interest largely from the perspective of the self-organization behavior of ants, such as how ants regulate traffic^9–11^. Trunk trails have also been studied from the perspective of their temporal and spatial dynamics as well as their energetic value in terms of efforts expended and resources obtained^12,13^. Yet, studies have not investigated how utilizing the same trails day after day impacts the exposure of ants to parasitism. Moreover, studies on ant foraging have largely occurred in a laboratory setting, and of the work that took place in the field, most studies relied on human observation or manipulated the environment in some way (see references in Supplementary Table S1). An ant species that forages collectively and predictably in time and space would be useful to assess the relationship between trail behavior and disease risk.

A potential system is the carpenter ant *Camponotus rufipes* in southeastern Brazil, which forms trunk trails lasting for multiple months^14,15^. Colonies of this ant were recorded as having a chronic infection by the fungal parasite *Ophiocordyceps camponoti-rufipides* across 20 months^16,17^. This fungus manipulates foragers to leave the nest and die biting the underside of a leaf ^17,18^. To complete its lifecycle, the fungus must grow out of the ant cadaver and form a fruiting body that releases spores onto the ground below that will infect other ants^18^. Cadavers are found attached to leaves surrounding the ant nest^17^. The chronic nature of infection at the colony level means the spores of the pathogen are continuously in the environment from the perspective of the foragers. The spores are curved and large (80-95 microns^16^) implying they do not travel far and land on the nearby trails once released from ant cadavers that hang above trails. Spores germinate to produce infectious secondary spores on hairs (capilliconidia) which attach to ants as they walk over them^19^. Thus, infection does not require a spore to hit an ant as it walks on a trail below a cadaver. Instead, the trail substrate itself serves as the source of contamination.

Foragers of the carpenter ant *C. rufipes* mostly collect nectar from hemipteran secretions and extrafloral sources^14,20^. The exploitation of a stable resource suggests that all foragers will emerge to walk directly to the food source, utilizing trails near the colony entrance as a highway^15^. Evidence from other systems demonstrate trunk trails as well organized for traffic flow^10^. Traffic is bi-directional on the trunk trails of *C. rufipes*. Thus, we expect an even mixture of inbound and outbound ants as this is hypothesized to increase flow^21^. If colonies can regulate the number of foragers on the trail to create a steady flow, we expect forager speed to remain approximately constant throughout the foraging period as foragers are not limited by the density of ants on the trail. Lastly, we are interested to see how the individual walking behavior observed influences the likelihood of an ant encountering an infectious spore.

We set out to study trails of seven *C. rufipes* colonies in their undisturbed rainforest habitat with both food sources and pathogens occurring at natural levels. We devised a system of recording trails using infrared lights and modified cameras to contend with the nocturnal foraging of this species. To overcome observer bias and ensure a larger body of data from which patterns may emerge, we used machine learning to automate ant tracking. This provided us with a powerful dataset from which the movement pattern of ants throughout a foraging period can be examined. We then characterized the forager trajectories on speed, straightness, and direction. Based on these measurements, we were able to classify ants on the trail into behavioral groups. Using an agent-based model based on our data, we suggest a mechanism for the maintenance of disease in this system.

## Methods

### Study site

Fieldwork took place at the Research Station of Mata do Paraíso, Universidade Federal de Viçosa, Minas Gerais, Southeast Brazil (20°48’08 S 42°52’31 W) between 10 and 25 January 2017. The carpenter ant *Camponotus rufipes* is abundant in this area, forming trails lasting multiple months^14,15^. Trails of *C. rufipes* are typically found on ‘bridges’ composed of woody debris, lianas and tree branches and are rarely directly on the forest floor^15^. Ants forage at night and activity peaks in the early evening^15^.

### Trail filming

Trails from seven different *C. rufipes* nests were filmed between 10 and 25 January 2017. Nests were selected based on their location and structure. Only nests found above the ground with nest material clearly visible were used. Trails were filmed before a branching point from the main trail so that ants were filmed coming directly from or towards the nest. In the case where multiple trails came from one nest, the busiest trails were selected. The width of the branches filmed ranged from 0.8 cm to 7 cm (mean ± standard deviation; 2.97 cm ± 2.53) and the length of the area filmed for all branches was approximately 15 cm.

GoPro cameras (model: HERO 3+, GoPro, Inc., San Mateo, USA) with a modified infrared filter (RageCams.com, Michigan, USA) were used for filming. Stakes were placed 30 centimeters from the trails and 30 cm medium trigger clamps (DWHT83140, DeWalt, Towson, USA) were attached to the stakes. Cameras were attached to clamps so that cameras were approximately 30 centimeters above the trails looking down at the ants walking on the trails (Supplementary Fig. S1). An additional camera was placed on the stake, looking sideways at the ants, to allow another perspective for behavioral analysis. Filming lasted from 19:30 to 00:00 for 4-7 nights for each trail (Supplementary Table S2). Timing of filming was based on previous work showing activity begins around 19:30 and peaks around 21:00^15^. Infrared lights (IR30, CMVision, Houston, USA) were connected 12-Volt 7Ah batteries (UP1270, UniPower, São Paulo, Brazil) to allow illumination of the trail without disturbing the behavior of the ants. The camera batteries lasted for approximately 1.5 hours, so the battery was changed once in the middle of a filming period. Slight adjustments in where the trail was positioned in the video view would sometimes occur at this time. Figure 1a shows an example image of a trail filmed and images of the remaining trails filmed are found in Supplementary Figure S2.

**Figure 1.**
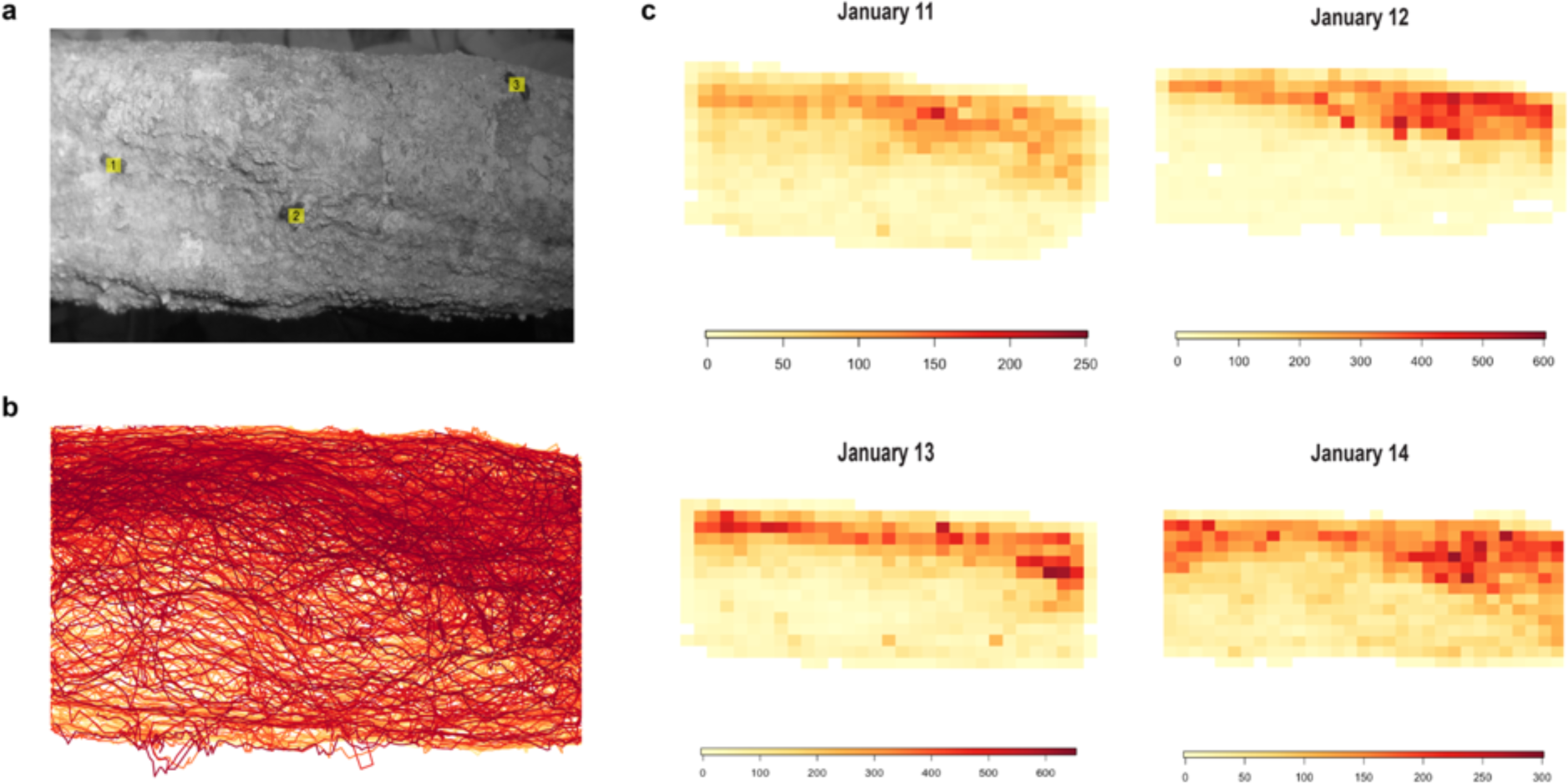
Trail image, trajectory overlay, and collective movement pattern. **(a)** Example image of a trail filmed taken from GoPro footage from colony MP1. Ants are labeled with identification numbers. **(b)** All of the trajectories from a single night of footage (January 14) at colony MP1. Each line across the trail represents a different ant, with the different colors distinguishing between different ant tracks. **(c)** The trail space from (a) was divided into a grid with each square representing approximately 1cm^2^. The number of times an ant walks into a square of the grid was calculated and the darker colors represent areas of the trail that ants walked over more. Each heatmap represents a different date (January 11 through January 14) from approximately the middle of the night to control for differences in the timing of filming. Different scales were used for each night, due to variance in the number of ants that walked across the trail.

### Automated ant tracking

A total of 78 hours and 56 minutes of video were recorded for seven colonies across four nights (Supplementary Table S2). We developed a machine learning approach to process and analyze these videos using a deep learning based segmentation model that identified ants as they came onto the screen and tracked them as they moved across the screen.

Our automatic ant tracking method contains two main processes: (1) detecting ants in each image frame of all videos, and (2) building ant trajectories for every video based on the detected ants. Commonly, deep learning schemes require a large amount of labeled ground truth data for model training. Since our dataset is quite large (> 8 million image frames), we aimed to generate sufficient labeled data for training our deep learning model without incurring excessive human labeling effort. Also due to the large size of our dataset, common active learning based sample selection methods (e.g.^22^) are not efficient. The goal of ant detection is to build ant movement trajectories and since ant trajectories normally span multiple consecutive frames in videos, detected ant positions in earlier frames assist with ant detection in later consecutive frames. That is, while ant detection forms a basis for building ant trajectories, trajectories of detected ants may also help ant detection. Hence, we designed our trajectory building procedure such that it not only can track detected ants but also can provide cues to indicate where (which frames and locations) there might be inconsistencies in ant trajectories and difficult scenarios for ant detection (e.g. densely clustered ants). We used such cues to select difficult cases from the frames for labeling to improve the deep learning detection model as well as the ant detection results. Therefore, our detection-tracking method consists of two rounds (with the second round improving the detection and tracking results of the first round), and each round performs two major steps, ant detection and trajectory building, as described below.

(1) Ant detection. This aims to detect ants in all the frames of the videos. We applied a novel object detection and segmentation model, Mask R-CNN ^23^, to automatically detect ants in every frame.

(2) Ant trajectory building. Given the detected ants in each frame, the next step is to form ant trajectories that connect detected ants frame-by-frame in videos. We formulated this ant trajectory building problem as a *transportation problem*, that is, between every two consecutive frames in each video, we find an optimal transportation (for ants) that corresponds to real movement of ants. In this transportation formulation, each detected ant in frame *K* can be viewed as a ‘supplier’ and each detected ant in frame *K+1* can be viewed as a ‘receiver’. The dissimilarity (based on spatial distance and appearance difference) between ants in two consecutive frames is a measure of how much ‘cost’ it would take to transport (move) one ant in frame *K* to another in frame *K+1*. The objective is to transport detected ants (as many as possible) in frame *K* to frame *K+1* with the minimum total cost. Optimal transportation based tracking methods are known to be effective for tracking sets of moving and changing objects in image sequences^24,25^.

In the first round, we randomly selected frames to label as training data. This allowed us to quickly and unbiasedly obtain data samples for training a decent detection model. We then applied the trained model to all of the frames to produce ant detection results. Next, we conducted trajectory building on detected ants to form the ant trajectories. Besides tracking ant movement, our trajectory building procedure in the first round also provided cues for identifying inconsistencies in ant trajectories and difficult cases in the frames for ant detection. In the second round, we applied training data selection to those difficult cases to find additional frames for labeling, and the enlarged training dataset thus obtained was used to re-train the Mask R-CNN detection model. The re-trained detection model was then applied to all the frames to produce the final ant detection results, which were used to build the final ant trajectories in the videos.

To identify difficult cases for additional training data selection, we used the following set of measures to capture possible errors in ant detection and trajectory results. (i) Ant speed: At a place where ants usually do not move very fast but a fast movement is suggested by the optimal transportation solution, this instance might indicate an error in ant detection. (ii) Missing ants in the middle part of a tree branch: When the optimal transportation solution does not find a corresponding ant instance in the next frame in the interior section of a tree branch, it might suggest a missing data point in ant detection. (iii) Ant identification (ID) switching: Each detected ant was assigned an ID number; when multiple ants are seen at spatially close interaction and slight changes on the dissimilarity scores among these ants give largely different solutions for the optimal transportation problem, this might suggest an ant ID switch error. Based on these observations and measures, our trajectory building process can help identify difficult detection and tracking cases for additional training data selection to improve model performance.

Overall, our automatic ant detection and tracking method extracted the *x* and *y* coordinates in pixels of detected ants in every frame and assigned each ant an identification number (Fig. 1a; Supplementary Video S1). Ant identification numbers were used to form ant trajectories used in further analysis.

### Error assessment

To assess the accuracy of the computer model, we watched a subset of videos and determined the error rate. GoPro cameras automatically divide footage into 26-minute-long videos, so one night of footage at a single trail has 6 to 10 videos. This provides a way of checking the accuracy of the computer tracking at random points throughout a night. We first error checked videos from the middle of the night (when the trails should be busiest) to determine if the data from that colony was high enough quality to use in our analysis. If the error rate was sufficiently low, we continued to error check all videos and nights for that colony. To error check, we counted the number of ant trajectories with errors out of the first 15-30 tracked ants. The number of ant trajectories checked varied because videos from early in the foraging period sometimes had fewer ants.

To ensure consistency in the type of ant trajectories that were analyzed, trajectories beginning in the middle of the field of video view were removed. This created uniformity between all colonies and nights in the type of ants that were compared as it focused on the ants that made it from one end of the trail to the other completely in the view of the video.

### Trajectory analysis

We used R version 3.4.4 and RStudio version 1.1.447 for all analyses^26,27^. Ant location data was frame-by-frame, so we used the native frame rate of the cameras (29.97 or 25 frames per second; the default setting of the cameras varied) to convert the time in frames to seconds and then used the start times of each video to convert it to real time (Supplementary Table S2). To convert ant location data from pixels to centimeters, we placed a ruler in each video to determine the conversion factor (Supplementary Fig. 2).

To determine how individual ants were moving, we calculated the following variables: average speed, overall direction, time on the trail, and straightness. Average speed was taken as the total distance an ant travels while in the video over the time it takes for them to travel that distance. Overall direction was whether the ant headed away from or towards the nest which we determined based on where the ant entered and exited the video view. A variety of measures are used to determine the straightness or tortuosity of an animal’s movement path ^28,29^. Ant movement on trunk trails is expected to move in an oriented direction, and not be a random search path, thus we used the simplest measure, the straightness index ^29^. The straightness index (ST) is a ratio between the net displacement and total path length:

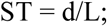

where d = the distance between the beginning and end of the path and L = total path length

### Agent-based model

To assess the influence of foraging style on disease risk, we developed an agent-based model in NetLogo 6.0.2^30^ based on the walking style of the ants in our videos (full details in Supplementary Materials). This model tested how walking straight influences the hypothetical number of spores an ant picks up. Spore density varied from 10% to 100% of the patches in the environment covered in spores. We varied spore density in 10% increments, leading to 10 different spore density conditions. The straightness of an ant varied from 0 to 1 in 0.01 increments, leading to 101 different straightness scores. The model was run 30 times for each combination of parameters (1010 total combinations) leading to a total of 30300 runs.

### Statistical analysis

A linear mixed-effects models fit was used to assess whether the speed of ant changes over a foraging period. The model was generated using the lmer function in the R package’ lme4’^31^, with speed as the fixed effect and colony and date as the random effects. The package ‘lmerTest’^32^ was used to generate p-values. We checked the plotted residuals to ensure homoscedasticity prior to utilizing the results of the model. We used linear regression to analyze the results of the agent-based model, with the straightness value as the predictor of proportion of spores picked up in the environment with a log transformation to control for skew.

### Data Availability

The original videos and data analyzed in this study will be accessible through ScholarSphere (https://scholarsphere.psu.edu/) upon publication of this study.

## Results

### Automated tracking performance

The automated tracking of ants in video frames resulted in 20,230,585 data points on ant movement. The model had two types of accuracy against which it can be judged, relative to a human. The first is species accuracy (detection accuracy) which is a measure of how well the model recognized the correct species of ant. The model correctly detected *C. rufipes* ants with an accuracy of 97.86%. The model picked up other insects or species of ants on the trail (false positive) or failed to detect a *C. rufipes* ant as it went across the trail 2.14% of the time.

The second accuracy measurement is tracking accuracy. The computer had to detect *C. rufipes* ants and follow them as they moved across the screen. If an ant moved in a straight line this required the computer to recognize and track that ant for about 4 seconds or 120 frames. The computer assigned identification numbers to individual ants to follow an ant as it travelled across the screen. The machine learning model sometimes made errors in doing this. The computer may switch identification numbers when ants walked too closely together (Supplementary Video S2). The average tracking accuracy for all colonies was 78.70%. The tracking accuracy was the lowest for MP2 (40.0%), MP11 (31.7%), and MP17 (50.6%). Identification number switches commonly happened in colonies MP2 and MP11. These trails were very thin and introduced more challenges in determining the trajectories of individual ants, so they were removed from further analysis. We have additionally removed MP17 as an obstruction in the trail led to ants departing from the branch and walking underneath leaves (Supplementary Video S3). Ants disappearing under leaf debris made it difficult to track an individual ant. We have made all videos and data available as we expect improved future machine learning models can make use of them.

The exclusion of these colonies brought the size of the dataset to 8,505,784 data points on ant movement from four colonies: MP1, MP6, MP10, and MP16. The large reduction of the number of data points from the elimination of 3 colonies can be attributed to the errors in these branches, where the density of individuals in congested areas lead to a false inflation of the number of ants and overall data points. The data points from the 4 included colonies represents the movement data for 64,499 ants. The average tracking accuracy of the remaining colonies was 81.39% (MP1: 72.0%; MP6: 82.1%; MP10: 77.2%; MP16: 92.1%). Most errors were due to an identification number switching to a different ant (8.28%). The high error rate for MP1 could be attributed to the darkness of the videos causing the model to miss part of an ant’s trajectory or failing to detect an ant in the dark areas of the trail. If we consider only the errors where a number is on a wrong ant or a number is not on an ant, the accuracy improves greatly (overall: 90.94%; MP1: 91.5%; MP6: 88.8%; MP10: 86.6%; MP16: 96.3%). We are mainly concerned with the direction and shape of trajectories, and the main error that impacts an individual ant’s trajectory is when ants switch to the wrong identification number, so the second calculation of accuracy rate is more reflective of this.

### Collective movement pattern

Most ants walk on the same area of the available trail space (Fig. 1). Ants often follow each other, walking across the same area (Supplementary Video S4). The trail usage pattern is consistent between nights (Fig. 1c). The mean speed of all ants from all colonies and nights was 5.19 cm/s ± 1.61 (standard deviation). The average speed of the colonies ranged from 4.74 cm/s to 5.62 cm/s and within colony variability in speed was similar between colonies (mean (cm/s) ± standard deviation; MP1: 4.99±1.69; MP6: 5.62±1.60; MP10: 4.88±1.53; MP16: 4.74±1.41). The results of the linear mixed effects model showed that ant speed decreases by 0.50 cm/s ± 0.07 (standard error) throughout the night (t_(96.45) =_ −7.12, p < 0.0001) (Supplementary Fig. S3).

### Individual movement pattern

Although most ants walked on the same area of the branch (Fig. 1b-c), there was a subset of ants that walked differently based on the straightness score (Fig. 2). Based on the behavioral analysis of videos, ants that had a straightness score of close to one walked straight across the trail as was expected (Fig. 2b; Supplementary Video S5). We found that 80.8% of ants had a straightness score from 0.75 to 1 (n = 50,813). We labelled these ants as ‘direct walkers’. Ants with an intermediate straightness score typically made it from one end of the trail to the other, but spent time wandering and covering more area of the trail (Fig. 2b; Supplementary Video S5). We labelled these ants as ‘wanderers’. They represent 13.0% of ants and had a straightness score from 0.25 to 0.75 (n = 8,194). By contrast, 6.2% of ants had a straightness score of less than 0.25 (n=3,869). These ants with a very low straightness score typically circled on the trail consequently entering and exiting on the same side of the video view (Fig. 2c; Supplementary Video S5). We labelled these ants as ‘circlers’.

**Figure 2.**
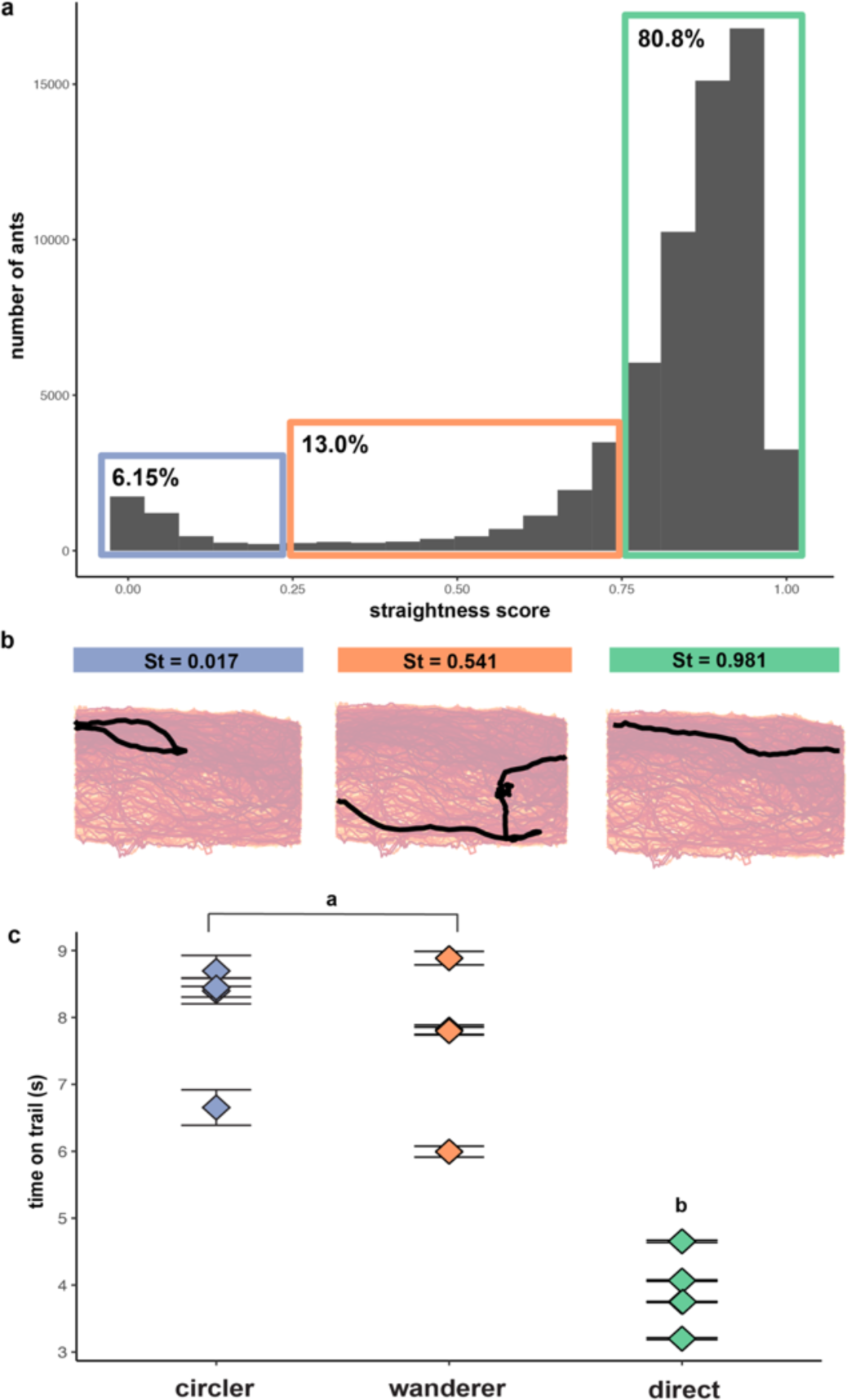
Different behavioral groups based on straightness score. **(a)** Histogram showing the distribution of straightness scores for all nights and colonies. **(b)** Example trajectories for a circler, wanderer and direct walker highlighted over all of the trajectories shown in Figure 1b. The straightness score (St) for that trajectory is included above. **(c)** Mean time spent moving across the trail in seconds for each different behavioral group and colony ± standard error of the mean. Different points within a behavioral group represent different colonies. Superscripts indicate groups as significantly different (p < 0.0001).

The wanderers and circlers constituted the minority of records (13% and 6.2% respectively). We observed these two behavioral phenotypes regardless of whether there were other ants in the area (Supplementary Video S5). These ants often stopped and groomed or antennated the trail or air (Supplementary Video S6). However, direct walkers were also observed stopping and grooming their antennae (Supplementary Video S7). There was a significant effect of straightness group on time spent on the trail for all three groups (Fig. 3e; one-way ANOVA; F_(2, 64495)_ = 14350, p < 0.0001). Post hoc comparisons using the Tukey Test indicates circlers did not spend more time on the trail (mean=8.1 seconds, SD=6.85) than wanderers (mean=7.58 seconds, SD=3.59), but both spent significantly more time on the trail than direct walkers (mean=3.88 seconds, SD=1.42).

**Figure 3.**
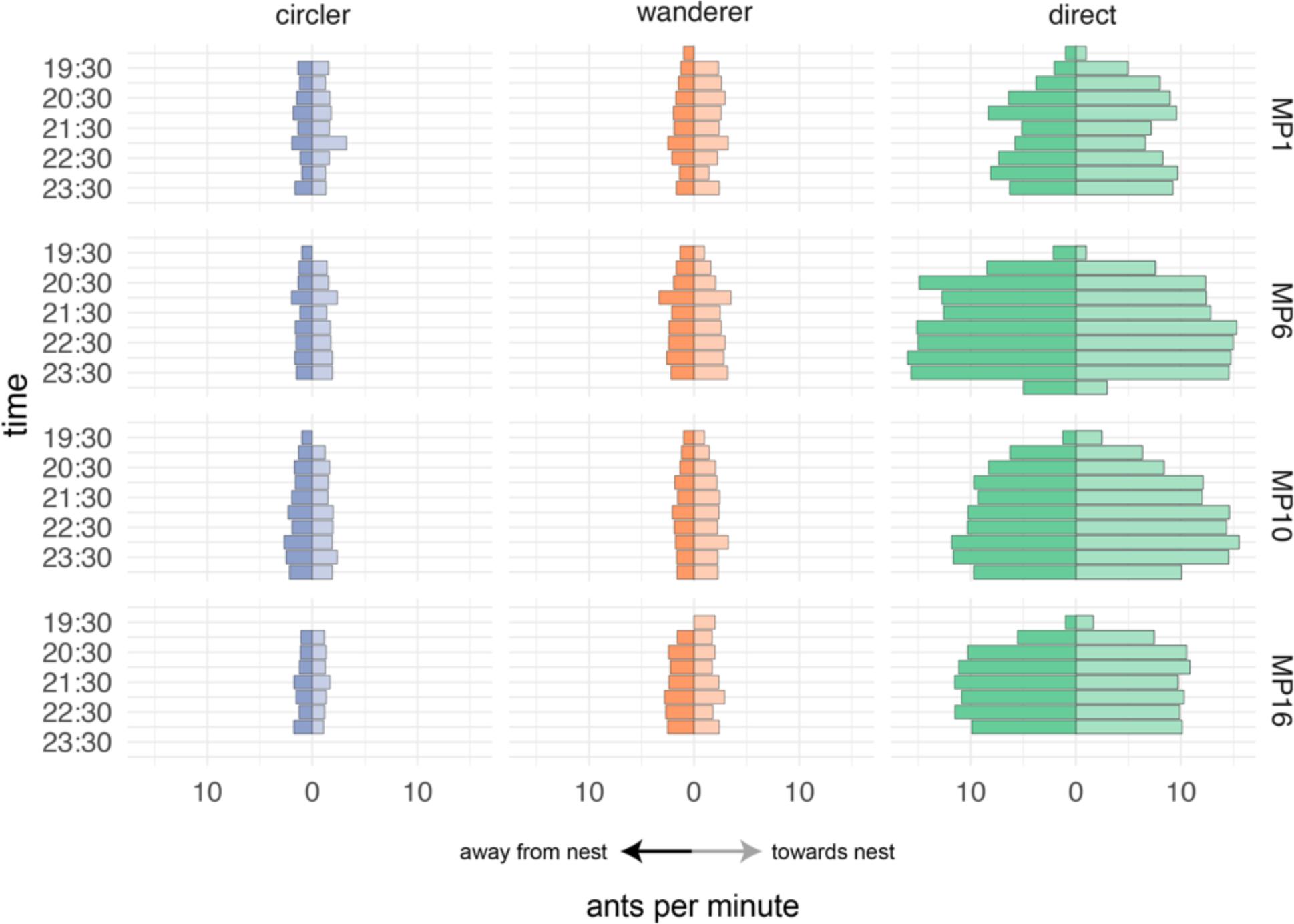
Distribution of different behavioral groups over time. Mean number of ants per minute in each behavioral group in a 30-minute period going either away from the nest or towards to the nest. Averaged across all nights for each colony. Right side numbers represent different colonies.

### Temporal movement pattern

The flow of all three groups of ants (direct walkers/wanderers/circlers) in and out of the nest was approximately the same throughout the night (Fig. 3). There is a large increase in the number of direct walkers on the trail throughout the night, while the number of wanderers and circlers throughout the night is relatively constant.

### Agent-based model

Based on the results obtained from our agent based model, walking straight significantly decreases the proportion of potential spores an ant picks up in an environment (linear regression: F_(1, 30298)_=5,458, p < 0.0001; Fig. 4a). The three different groups of foragers differed in the number of spores they pick up in the environment regardless of spore density (F_(2, 30297)_=21,208, p < 0.0001; Fig. 4b). Circlers pick up significantly more spores than wanderers and wanderers may pick up significantly more spores than direct walkers (Tukey Test; p < 0.0001).

**Figure 4.**
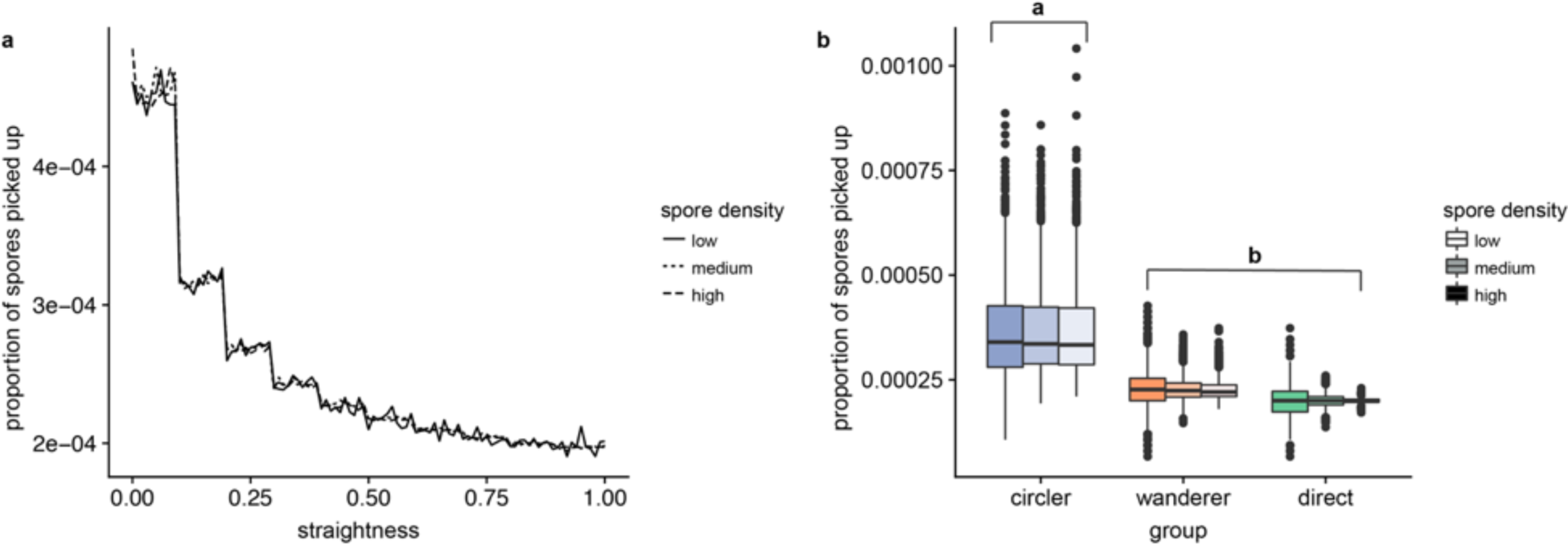
Risk of spore exposure for different behavioral groups. **(a)** Mean proportion of spores picked up as simulated ants in the agent-based model walk across the trail with different straightness scores **(b)** Data from (a), with straightness divided into behavioral groups. Superscripts indicate groups as significantly different (p < 0.001).

## Discussion

Our study utilized an unobtrusive filming set-up to record behavioral data on more than 64,000 ants moving in a rainforest at night in an area of high disease pressure. The study design facilitated the capture of natural ant behavior unaffected by either a laboratory environment or proximity to human observers. Combining this approach with computer vision techniques increases the scale at which we can study animal behavior. Using computer vision and deep learning we collected approximately 20 million ant movement data points from 80 hours of nighttime video. A previous study, using humans to score the positions of ants in each frame, required approximately 1,600 hours of human work to create a dataset of 6.9 million data points (Modlmeier et al., in review). Advances in camera technology improving our nighttime recording capabilities along with increased computing power allowing machine learning to identify individuals promotes research on natural animal behavior.

For our study on ant behavior in the context of disease transmission, the scale of this data detected higher level patterns likely unobservable with a less detailed dataset. Our data shows ants flowing in and out of the nest at approximately the same rate (Fig. 3). Work on harvester ants (*Pogonomyrmex barbatus*) has shown that the feedback from returning foragers stimulates inactive foragers to leave on a new trip^33^. Our even flow rate also validates work on Argentine ants (*Linepithma humile*) showing ants exiting and entering the nest at approximately the same rate in the summer^34^. Ant colonies operate through local interactions and without centralized control, so there is no authority controlling when ants leave and return to the nest^35,36^. The lack of centralized control combined with the even flow rate gives insight into the processes occurring within a nest, with returning foragers likely stimulating new foragers to leave the nest.

Given that there is no centralized plan for foraging, it is impressive that the same foraging pattern arises on different days (Fig. 1c). This consistent trail usage pattern, along with most ants walking straight across the trail (Fig. 2a) likely emerges from the use of a chemical trail, which this species of ant (*C. rufipes*) is known to use^37^. For ants to walk on the same area of the trail on different nights, the trail pheromone must either persist between foraging periods or foragers repeatedly reinforce the fastest route across the branch each night. While rare, we observed some ants on these trails during the daytime, and other studies have observed *C. rufipes* foraging during the day^20^. This could allow the trail to be reinforced around the clock. Alternatively, laboratory studies have demonstrated ants as preferentially selecting the shortest route to food^38,39^. The path that receives more pheromone will be reinforced quicker^40^. Thus, each night the portion of the trail that ants walk on fastest could reach a higher concentration of trail pheromone quicker, leading to the pattern observed.

The texture of the tree branch could also drive the space usage pattern, as substrate and landscape features impact ant locomotion^41,42^. Loreto et al. (2013) demonstrated, in the same population we studied, that *C. rufipes* foragers in this environment prefer to walk on woody debris because they walk faster on this material than on the forest floor (see Supplementary Video S8 for an example of how ants are impeded on the forest floor). The type of wood could also make a difference, with ants preferring to walk on areas of the trail that are least restrictive to their movement. Another pattern emerged through investigation of the straightness index of the ants. As expected, the straightness index of most ants was close to one (80.8%; n=50,813), indicating that they walked directly across the trail (Fig. 2). Ants may prefer on the path that deviates the least from their original direction of travel^43^. Straighter individual paths enhance information spread and increase the chance that an ant will find food^44,45^, perhaps making this pattern beneficial to the collective colony in resource acquisition.

Despite the dominance of ants walking straight across the trail, a proportion of foragers wandered before making it across the trail and another group circled the trail and headed back in their original direction (Supplementary Video S5). Other species of ants provide evidence of different roles within foragers, such as patrolling, trail maintenance, and defense. Patrollers in harvester ants are the first to leave the nest in a foraging period and determine which trails the colony will use that day^46^. However, if the subtypes in this study were patrollers, we would expect there to be more of them at the beginning of the night which is not the case (Fig. 3). The leaf cutting ant *Atta cepahlotes,* which also forms consistent trails, has a special class of foragers involved in trail maintenance^13,47^. Ants were observed carrying leaves (Supplementary Video S9), although this could be for nest material and not trail cleaning. Another role could be maintaining the pheromone trail. For example, *Atta sexdens* minims help with the pheromone trail instead of food transport^47^. Ants were observed dragging their gaster on the trail likely depositing trail pheromone (Supplementary Video S10). However, it seems unlikely wanderers and circlers were involved in maintaining the pheromone trail, as they should walk straight across the trail to ensure the pheromone trail was on the most direct path.

The subtypes could also be involved in defense. Wanderers and circlers spent more time on the trail (Fig. 2c) and were observed stopping and antennating (Supplementary Video S8). Smaller workers hitchhike on leaf fragments carried by larger workers in *Atta colombica* leaf-cutting ants, and this likely serves as a defense against parasitoid Phorid flies^48^. Flies, that could possibly be parasitoids, were observed closely following ants on the trail and in some cases appearing to land which may indicate laying eggs on the ants which later become endoparasitoids (Supplementary Video S11) although the prevalence of parasitoid flies attacking *C. rufipes* is unknown. We have observed adult ants infected by decapitating phorid flies in our study area (Supplementary Video S12).

Conversely, the forager variation in walking straight may not indicate different roles within the colony and instead demonstrate differences in response plasticity, as individuals may differ in their detection of the pheromone trail. Bumblebees (*Bombus terrestris*) vary in their antennal sensitivity to odors and different behavioral thresholds have been found for castes of the ant *Pheidole pallidula*^49,50^. Heterogeneity can be beneficial to the collective colony in tasks such as selecting a new nest^51^. In the context of trunk trail foraging, it could encourage exploration and increase the chance of discovery of high value food items. *Camponotus rufipes* typically uses trunk trails to exploit dependable food resources such as hemipteran secretions or extrafloral secretions. If the colony always remains on the trunk trails, they might deplete their dependable source of food and not have a suitable alternative. Argentine ants are able to adapt trails in response to resource availability^52^. We suggest wandering and circling as a mechanism for *C. rufipes* ants to similarly respond to changes in resource availability. We filmed only a small area of the foraging trails, providing a brief snapshot of an ant’s behavior. To know whether wanderers or circlers are more likely to wander from the trail and discover new food resources, one would need to follow individual ants during an entire foraging trip, which was beyond the scope of this study.

Following individual ants for their entire foraging trip would also clarify whether the straightness groups represent fixed behavioral groups or if they just demonstrate variation in individual behavior over time. Campos et al. (2016) studied the activity patterns of *Aphaenogaster sensilis* ants and found foraging trajectories to be descriptively similar with individual temporal activity patterns showing greater variation. In the context of our study, perhaps all ants engage in wandering or circling behavior on these trails, and it is related to their temporal activity pattern and not their behavioral role.

Regardless of whether these are fixed subtypes within the colony, variability in walking behavior could impact the maintenance of disease in this environment. Fungal infected cadavers surround these trunk trails, likely dropping spores directly onto the trails below^17^. It is not possible to quantify the abundance and distribution of micron sized spores on trails in a forest, but the long term tracking of cadaver abundance and the proximity to the trails implies spore presence on the foraging trails^17^. According to our simulations, walking in a straight line reduces a forager’s risk of picking up spores (Fig. 4). If all ants walked in exactly the same straight line, this could prevent the disease cycle from continuing, especially since the first ants would initially clear all of the spores off. Yet, this does not occur as the circlers and wanderers deviate from the straight path increasing their probability of picking up spores and maintaining a chronic infection of the colony.

If the risk of infection is larger for the circlers and wanderers (Fig. 4), why do these subgroups still exist? Social insects have members of the colony known as scouts that assist in discovering and recruiting the colony to new food sources^54–57^. The wandering and circling behavior observed in this study could reflect the individual ant’s role in food discovery, instead of food retrieval. Colonies with this variability in forager behavior are perhaps better able to obtain resources, improving their reproductive success and maintaining the diversity in forager behavior. Simultaneously, it allows persistence of the fungal parasite in the system, but the loss these ants is likely not enough to have a selective impact on the colony, as it is only a small percentage of foragers as suggested by Loreto et al. (2014). In the harvester ant *Pogonomyrmex owyheei*, less than 10% of the colony foraged and it was usually less than 6% at one time^58^. Reproductive success was also hardly impacted when harvester ant foragers were restricted from foraging^59^ implying colonies usually gather more than enough food and fitness would likely not take a huge hit from the loss of a few foragers. Scharf et al (2017) additionally demonstrated that colony fitness (as measured through the number of reproductive individuals produced) remained unchanged from parasitic infection. In our system, relatively few foragers appear to be infected and killed by the parasite^17^. In addition, the density of spores in the trails could be very low, since the trail network occupies less than 2% of the nest surrounding area^15^. Understanding the direct relationship between ant cadavers (from where spores are shot) and the trails (where new hosts are found) would improve our agent-based model predictions and help to understand the importance of wanderers and circlers for colony disease risk management.

Although these behavioral subtypes are only a small proportion of the colony, that small proportion could be more than enough ants to sustain a parasitic fungus. Understanding how variation in behavior influences pathogen risk provides information on the factors that shape the distribution of animals in time and space. Computational techniques serve as a way to collect large datasets on animal behavior, where one can begin to unearth the complex interactions between an animal and its habitat.

## Acknowledgements

We thank the Department of Forest Engineering at the Federal University of Viçosa for allowing us to perform this study at the Research Station of Mata do Pariso and Dr. Simon Elliot for hosting us in his laboratory. We are grateful to Charissa de Bekker who helped capture the behavior in Video S11. This work was supported in part by National Science Foundation Grants IOS-1558062 and EEID 1414296 to D.P.H, NSF CCF-1617735 to D.Z.C., and NIH Grant R01 GM116927-02 to D.P.H. and D.Z.C.

## Author Contributions Statement

D.P.H., R.G.L, and N.I. conceived and designed the study. N.I. and C.K. performed the field work with technical input from R.GL․ Y.Z. and D.Z.C. created the computer model and processed the data. N.I. analyzed the data and wrote the manuscript with guidance from D.P.H․ All authors reviewed the manuscript.

## Additional Information

### Competing Interests

The authors declare no competing interests.

